# Cyanobacterial growth and morphology are influenced by carboxysome positioning and temperature

**DOI:** 10.1101/2020.06.01.127845

**Authors:** Rees Rillema, Joshua S. MacCready, Anthony G. Vecchiarelli

## Abstract

Cyanobacteria are the prokaryotic group of phytoplankton responsible for a significant fraction of global CO_2_ fixation. Like plants, cyanobacteria use the enzyme Ribulose 1,5-bisphosphate Carboxylase/Oxidase (RuBisCO) to fix CO_2_ into organic carbon molecules via the Calvin-Benson-Bassham cycle. Unlike plants, cyanobacteria evolved a carbon concentrating organelle called the carboxysome - a proteinaceous compartment that encapsulates and concentrates RuBisCO along with its CO_2_ substrate. In the rod-shaped cyanobacterium *Synechococcus elongatus* PCC7942, we recently identified the McdAB system responsible for uniformly distributing carboxysomes along the cell length. It remains unknown what role carboxysome positioning plays with respect to cellular physiology. Here, we show for the first time that a failure to distribute carboxysomes leads to a temperature-dependent decrease in cell growth rate, cell division arrest, cell elongation, asymmetric cell division, and a significant reduction in cellular levels of RuBisCO. Unexpectedly, we also report that even wild-type *S. elongatus* undergoes filamentous growth at the cool, but environmentally-relevant, growth temperature of 20°C. The findings suggest that carboxysome positioning by the McdAB system functions to maintain the carbon-fixation efficiency of RuBisCO by preventing carboxysome aggregation, which is particularly important at temperatures where rod-shaped cyanobacteria adopt a filamentous morphology.

**IMPORTANCE:** Photosynthetic cyanobacteria are responsible for almost half of global CO_2_ fixation. Due to eutrophication, rising temperatures, and increasing atmospheric CO_2_ concentrations, cyanobacteria have recently gained notoriety for their ability to form massive blooms in both freshwater and marine ecosystems across the globe. Like plants, cyanobacteria use the most abundant enzyme on Earth, RuBisCO, to provide the sole source of organic carbon required for its photosynthetic growth. Unlike plants, cyanobacteria have evolved a carbon-concentrating organelle called the carboxysome that encapsulates and concentrates RuBisCO with its CO_2_ substrate to significantly increase carbon-fixation efficiency and cell growth. We recently identified the positioning system that distributes carboxysomes in cyanobacteria. However, the physiological consequence of carboxysome mispositioning in the absence of this distribution system remains unknown. Here we find that carboxysome mispositioning triggers temperature-dependent changes in cell growth and morphology as well as a significant reduction in cellular levels of RuBisCO.

## INTRODUCTION

Cyanobacteria represent a phylum of diverse prokaryotic organisms where many fundamental biological processes have remained largely understudied. As the evolutionary ancestor of algae and plant chloroplasts, all cyanobacteria perform oxygenic photosynthesis and fix carbon dioxide through the Calvin-Benson-Bassham cycle. Unlike chloroplasts, cyanobacteria encapsulate their RuBisCO and carbonic anhydrase within large (~150 nm) selectively permeable protein-based organelles called carboxysomes. This mechanism generates an environment around RuBisCO that is significantly enriched in CO_2_, which increases the carboxylation activity of RuBisCO, while simultaneously reducing photorespiration (1).

In the model rod-shaped cyanobacterium *Synechococcus elongatus* PCC 7942 (hereafter *S. elongatus*), carboxysomes were found to be uniformly distributed down the length of individual cells (2). This equidistant positioning supports equal inheritance of carboxysomes following cell division and maximum diffusion of substrates and products across the carboxysome shell. Carboxysomes are essential for the growth and survival of all cyanobacteria and are responsible for ~35% of global carbon-fixation through atmospheric CO_2_ assimilation (Dworkin, 2006; Kerfeld and Melnicki, 2016). However, it remains unknown how the subcellular organization of carboxysomes influences cyanobacterial physiology; a question of considerable ecological, evolutionary, and biotechnological importance.

Savage *et. al.* were first to report that a ParA-type ATPase (hereafter McdA - <u>M</u>aintenance of <u>c</u>arboxysome <u>d</u>istribution <u>A</u>) is required for positioning carboxysomes in *S. elongatus* (2). ParA family members have well established roles in the segregation of bacterial chromosomes and plasmids (4, 5). Less studied are ParA family members shown to be required in the positioning of diverse protein complexes, such as those involved in secretion (6, 7), chemotaxis (8–10), conjugation (11), cell division (12, 13), and cell motility (14, 15), as well as bacterial microcompartments (BMCs), such as the carboxysome (2, 16). We recently identified a small novel protein, McdB, responsible for generating dynamic McdA gradients on the nucleoid (16). McdB colocalizes and directly interacts with carboxysomes and removes McdA from the nucleoid in their vicinity. We found that carboxysomes use a Brownian-ratchet mechanism whereby McdB-bound carboxysome motion occurs in a directed and persistent manner toward increased concentrations of McdA on the nucleoid. We also recently found that the McdAB system is widespread among β-cyanobacteria (17); an incredibly diverse and widely distributed phylum of bacteria that display complex morphologies (18). Together, the data suggest that the equidistant positioning of carbon-fixing carboxysomes in cyanobacteria is important, but the physiological consequences of carboxysome mispositioning in the absence of the McdAB system remain unclear.

When Savage *et. al.* first identified the McdA requirement for carboxysome positioning, only a minor decrease in CO_2_ fixation was found in a Δ*mcdA* strain, but growth at 30°C under ambient CO_2_ was the only condition studied (2). Rising sea surface temperatures and coastal eutrophication in marine ecosystems are pervasive effects of climate change (19). Rising temperatures increase enzyme kinetics (20), leading to higher rates of protein synthesis, RuBisCO activity, and photosynthesis (21). Temperature therefore directly influences cyanobacterial growth and metabolic rates (22, 23).

Here, we show for the first time that a failure to distribute carboxysomes leads to temperature-dependent decreases in growth rate, cell division arrest, cell elongation, asymmetric cell division, and a significant reduction in cellular levels of RuBisCO. We also report that, unexpectedly, wild-type *S. elongatus* undergoes filamentous growth at 20°C; an environmentally relevant growth temperature for *S. elongatus* not commonly used in the lab. We propose that carboxysome positioning by the McdAB system functions as part of an autotrophic growth strategy that maintains the carbon-fixation efficiency of RuBisCO by preventing carboxysome aggregation. In the absence of carboxysome positioning or when cells are grown at cooler temperatures, we propose that cell elongation and asymmetric cell division are responses to organic carbon limitation, due to the decreased enzymatic activity of RuBisCO under these conditions.

## RESULTS

We performed *in vivo* microscopy to determine how carboxysome organization was altered in *mcdA*, *mcdB*, and *mcdAB* deletion strains compared to wild-type *S. elongatus*. In *S. elongatus*, each RuBisCO enzyme is composed of eight <u>L</u>arge (Rbc<u>L</u>) and eight <u>S</u>mall (Rbc<u>S</u>) subunits to form RbcL_8_S_8_ (24). Therefore, to image carboxysomes, the fluorescent protein mTQ (monomeric Turquoise2) was fused to the C-terminus of RbcS to make RbcS-mTQ (see Methods for strain construction details). *RbcS-mTQ* was expressed using a second copy of its native promoter (inserted at neutral site 1) in addition to wild-type *rbcS* at its native locus in wild-type cells as well as in the three deletion strains – Δ*mcdA*, Δ*mcdB*, and Δ*mcdAB*. The presence of RbcS-mTQ did not alter growth rate **(Supplemental Figure S1)**. Carboxysomes of *S. elongatus* encapsulate between 800 to 1,500 RuBisCO enzymes (25). Therefore, RbcS-mTQ provided a bright and high-contrast marker for quantifying the subcellular organization of carboxysomes, as previously shown (16). We also performed Phase Contrast imaging to monitor for potential changes in cell morphology, and Chlorophyll fluorescence imaging to verify that cells treated in our analyses were photosynthetically active and therefore viable.

In the lab, *S. elongatus* is typically grown at 32°C; with a doubling time on the order of several hours depending on growth conditions (temperature, light, and CO_2_ availability). This was the temperature used in our previous study that identified the McdAB carboxysome positioning system (16). However, *S. elongatus* was recently shown to grow faster at 40°C (26, 27). Therefore, in anticipation of a growth defect in the absence of carboxysome positioning in our Mcd system mutants, we began our studies by growing cultures at 40°C in constant light under ambient (0.04%) or high (2%) CO_2_ concentrations.

### Carboxysome mispositioning correlates with minor defects in cellular physiology at 40°C

We first verified that carboxysomes are organized by the McdAB system at 40°C, as we found previously at 32°C (16). We examined carboxysome positioning under high CO_2_ conditions (2% CO_2_) to maximize growth. We found that RbcS-mTQ labeled carboxysomes were equally spaced down the long axis of wild-type *S. elongatus* cells **(Figure 1A)**. The Δ*mcdA*, Δ*mcdB*, and Δ*mcdAB* mutants lost this uniform positioning of carboxysomes **(Figure 1B-D)**; as observed previously at 32°C (16).

**Figure 1.**
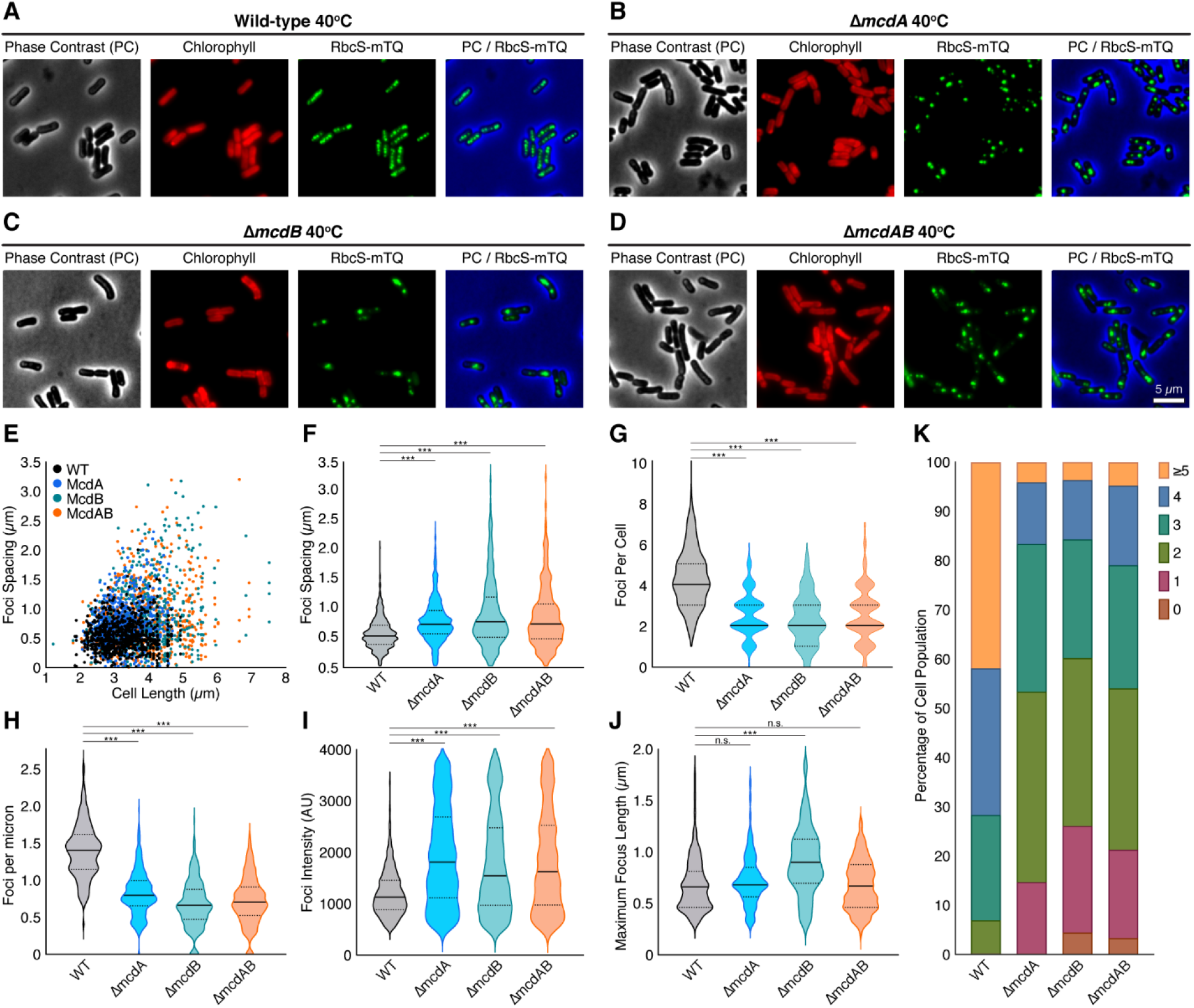
McdAB system mutants display fewer and mispositioned carboxysome aggregates at 40°C. (**A-D**) Microscopy images of the specified cell strains grown at 40°C in 2% CO_2_. Phase contrast (black), chlorophyll (red), and carboxysomes (green). Phase Contrast (PC) channel is blue in the merge for better contrast. (**E**) Spacing between carboxysome foci as a function of cell length. (**F**) Spacing between carboxysome foci in the same cell. (**G**) Carboxysome foci number per cell. (**H**) Number of carboxysome foci per unit cell length for each strain. (**I**) Carboxysome peak foci intensity for each cell strain. AU = Arbitrary Units. (**J**) Quantification of the maximum length across carboxysome foci. (**F-J**) Solid bars represent the median and dashed lines demarcate the 95% confidence interval. Statistical significance was based on a nonparametric Mann-Whitney test. *** = P < 0.001, ** = P < 0.01, * = P < 0.05, n.s. = not significant. (**K**) Population percentages of cells with the specified number of carboxysome foci. n ≥ 1000 carboxysomes from 440 cells of each strain.

We compared the nearest-neighbor spacing of carboxysome foci as a function of cell length **(Figure 1E)**. Wild-type cells showed uniform carboxysome spacing (0.55 ± 0.20 μm) regardless of cell length. All three mutants, on the other hand, displayed a gradual increase in the nearest-neighbor spacing of carboxysome foci as cell length increased. Both the median and variability in spacing were greater in the mutants compared to wild-type **(Figure 1F)**. The increased spacing resulted in fewer carboxysome foci per cell **(Figure 1G)**, and within single cells, there were fewer carboxysome foci per unit cell length **(Figure 1H)**. When quantifying the fluorescence intensity **(Figure 1I)** and size of carboxysome foci **(Figure 1J)**, it became apparent that the increased spacing in all three mutant populations was not likely a result of fewer carboxysomes being assembled, but rather, carboxysomes were coalescing into massive aggregates. Indeed, when comparing carboxysome foci number across cell populations, we find that ~ 95 % of wild-type cells (n = 440) had three or more foci, whereas ~ 80 % of all mutant populations had three or fewer foci **(Figure 1K)**. Roughly 20% of cells from all three mutant populations had a single carboxysome aggregate, whereas wild-type cells never had less than two foci. The 40°C data shows that the McdAB system not only equally spaces carboxysomes down the cell length, it also serves as an anti-aggregator; preventing carboxysomes from coalescing.

We then asked if carboxysome mispositioning affected cell physiology at 40°C. Wild-type *S. elongatus* cells were 3.1 ± 0.5 microns in length **(Figure 2A)** and 1.25 ± 0.05 microns in width **(Figure 2B)**. We found that *ΔmcdA* cells were of a similar length and width compared to that of wild-type **(Figure 2AB)**. The *ΔmcdB* and *ΔmcdAB* cells had a slightly longer median cell length, but a significantly wider distribution **(Figure 2A)**. These mutants were also thinner compared to wild type **(Figure 2B)**. The presence of both longer and shorter cells suggested asymmetric cell division events. We quantified the frequency of symmetric (mid-cell) versus asymmetric (non-mid-cell) division events and found that asymmetric division was exclusive to the *ΔmcdB* and *ΔmcdAB* populations **(Figure 2C)**. The data show that while both McdA and McdB are required for positioning carboxysomes, the loss of McdB elicits an asymmetric division phenotype at 40°C.

**Figure 2.**
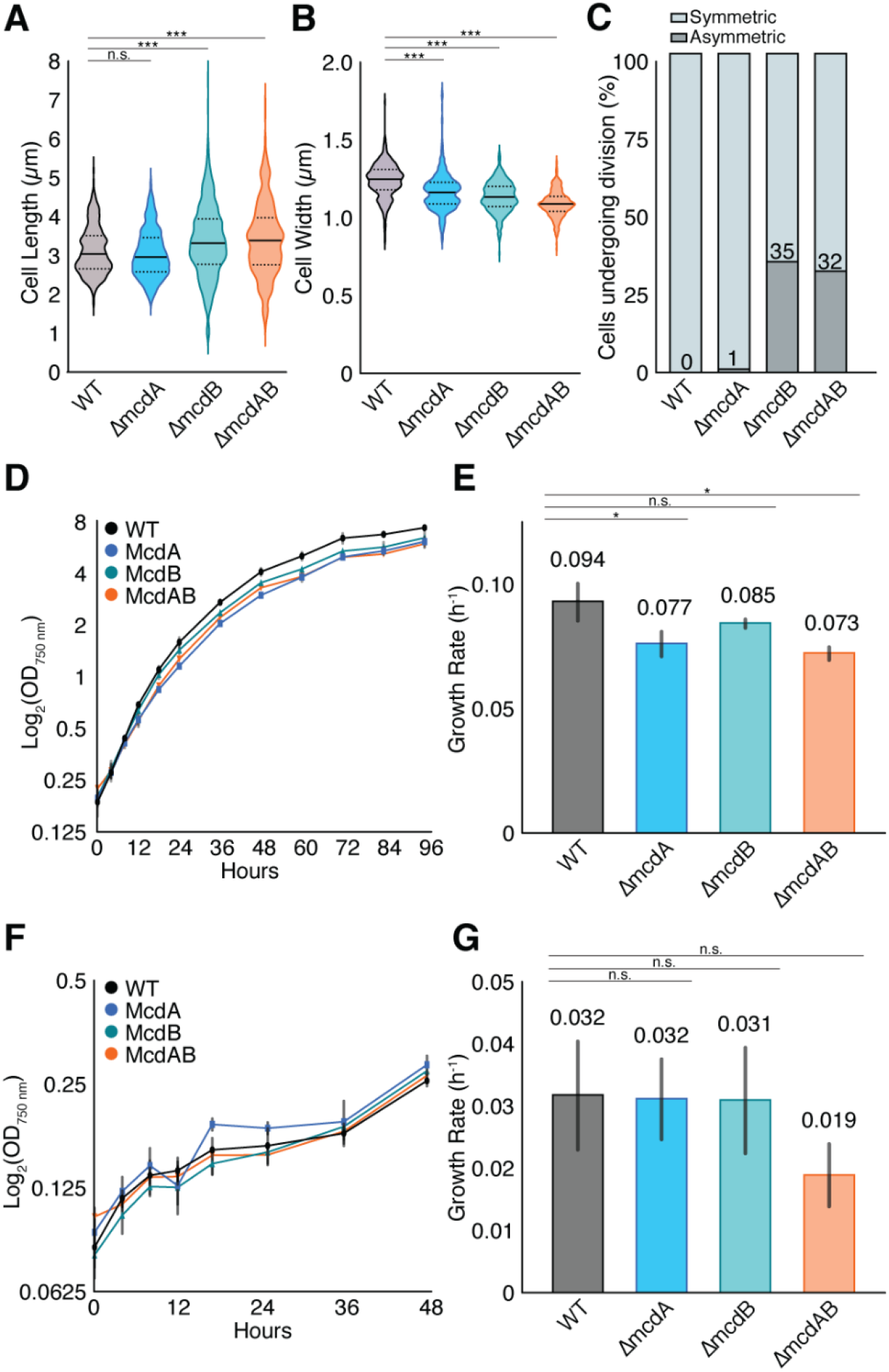
Carboxysome mispositioning correlates with subtle changes in cell physiology at 40°C. (**A**) Cell lengths and (**B**) widths of WT population compared to McdAB system mutants. Cells were grown at in 2% CO_2_. n = 440 cells of each strain. Statistical significance was based on a nonparametric Mann-Whitney test. *** = P < 0.001, ** = P < 0.01, * = P < 0.05, n.s. = not significant. (**C**) Percentage of cells undergoing symmetric (mid-cell; light grey bar) or asymmetric (non-mid-cell; dark grey bar) division. (**D**) Growth curves of all strains grown at 40°C in 2% CO_2_. (**E**) Quantification of mean exponential growth rate at 40°C in 2% CO_2_. (**F**) Growth curves of all strains grown at 40°C in 0.4% CO_2_. (**G**) Quantification of mean exponential growth rate at 40°C in 0.4% CO_2_. (**D-G**) Error bars represent the standard deviation from three independent biological replicates. Statistical significance was based on an unpaired t-test. *** = P < 0.001, ** = P < 0.01, * = P < 0.05, n.s. = not significant.

The changes in cell morphology suggested that, even with high CO_2_ levels, carboxysome mispositioning may alter cell growth. However, we found only modest reductions in growth rate for all three mutants **(Figure 2D-E)**. Even when grown slowly in ambient CO_2_ (0.04%), we did not observe significant differences in growth rate compared to wild-type **(Figure 2F-G)**. Overall, at 40°C, carboxysomes are mispositioned in all three Mcd mutants, with moderate increases in carboxysome spacing due to aggregation, which ultimately results in fewer carboxysome foci per cell. We found an asymmetric cell division phenotype exclusive to cell without McdB. But in all deletion strains, growth rate was not significantly slower than wild type, with or without CO_2_, at this temperature.

### Carboxysome mispositioning correlates with cell division arrest, cell elongation, and asymmetric cell division at 30°C

We continued our study at 30°C, a growth temperature closer to what we used to first identify the McdAB carboxysome positioning system (16). As we showed previously, and similar to our 40°C data, wild-type *S. elongatus* cells have equally spaced carboxysomes **(Figure 3A)**, while in all three mutants, carboxysomes were mispositioned **(Figure 3B-D)**. Wild-type cells displayed the same carboxysome spacing distance (0.50 ± 0.20 mm) regardless of cell length, while all three mutants had increased carboxysome spacing, and variability in spacing, as cell length increased **(Figure 3E-F)**. Intriguingly, we found that mutant cell lengths were significantly longer compared to wild-type. This cell elongation phenotype was more extreme in the Δ*mcdB* and Δ*mcdAB* mutants, resulting in more distantly spaced carboxysome foci **(Figure 3F)**. The data once again shows that McdB plays a currently unknown role in carboxysome function, separate from its role in positioning with McdA.

**Figure 3.**
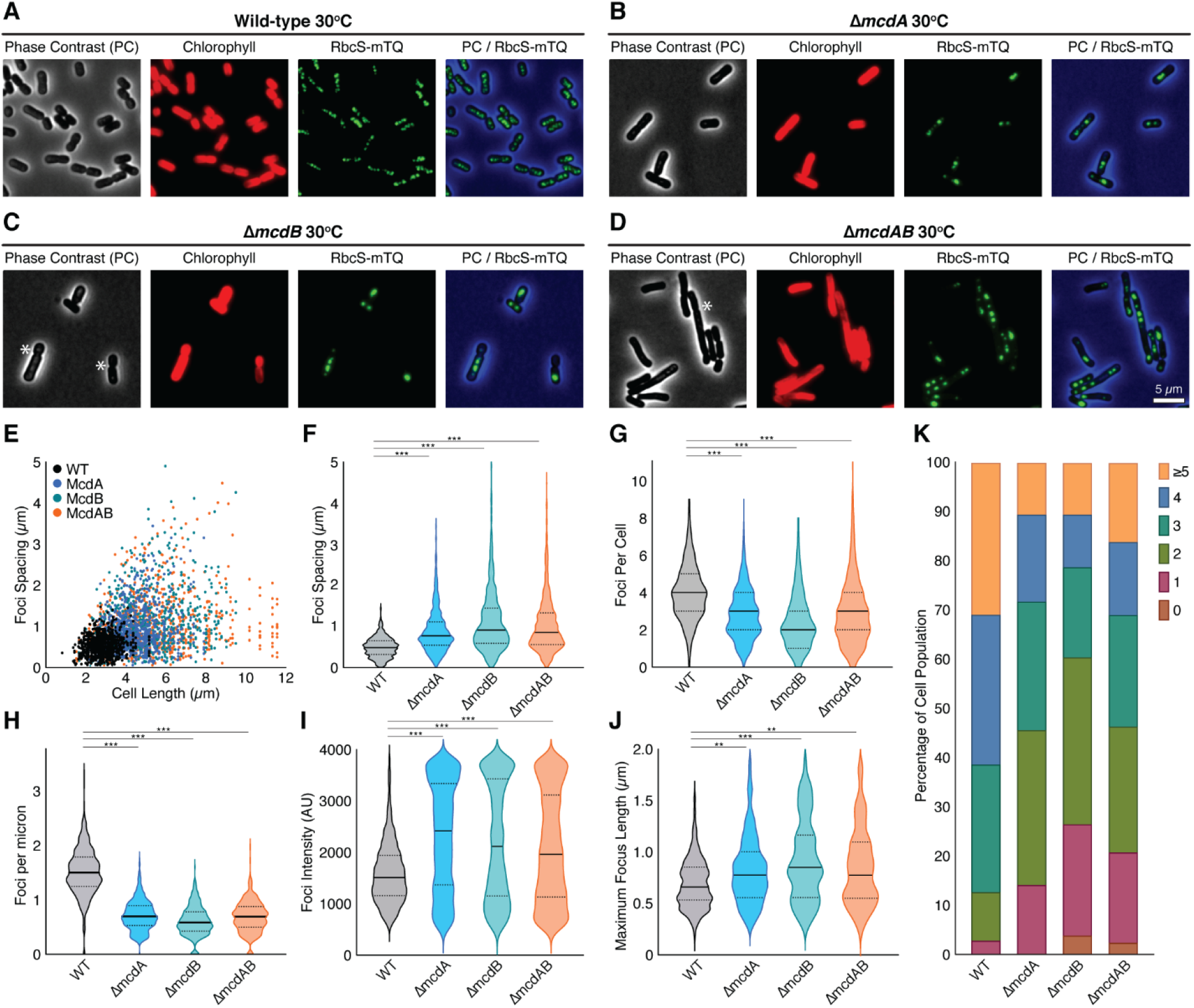
McdAB system mutants house few and mispositioned carboxysome aggregates at 30°C. (**A-D**) Microscopy images of the specified cell strains grown in 2% CO_2_. Phase contrast (black), chlorophyll (red), and carboxysomes (green). Phase Contrast (PC) channel is blue in the merge for better contrast. (**E**) Spacing between carboxysome foci as a function of cell length. (**F**) Spacing between carboxysome foci in the same cell. (**G**) Carboxysome foci number per cell. (**H**) Number of carboxysome foci per unit cell length for each strain. (**I**) Carboxysome peak foci intensity for each cell strain. Arbitrary Units (AU). (**J**) Quantification of the maximum length across carboxysome foci. (**F-J**) Solid bars represent the median and dashed lines demarcate the 95% confidence interval. Statistical significance was based on a nonparametric Mann-Whitney test. *** = P < 0.001, ** = P < 0.01, * = P < 0.05, n.s. = not significant. (**K**) Population percentages of cells with the specified number of carboxysome foci. n ≥ 1000 carboxysomes from 440 cells of each strain.

The increased spacing resulted in fewer carboxysome foci per cell **(Figure 3G)**, and per unit cell length **(Figure 3H)**. Carboxysome foci in all three mutants were significantly larger than that of wild-type, suggesting aggregation **(Figure 3I-J)**. While ~ 90 % of wild-type cells (n = 486) had three or more foci, ~ 70 % of all mutant populations had three or fewer foci **(Figure 3K)**. Again, ~ 20 % of cells in all three mutant populations contained a single carboxysome aggregate. This is a striking reduction in carboxysome foci when considering the cell elongation phenotype exclusive to the mutant cell lines.

As the carboxysome spacing data suggested, all three mutants had significantly longer cell lengths compared to wild-type when grown at 30°C **(Figure 4A)**. Median cell width was similar across all strains, however *ΔmcdB* and *ΔmcdAB* cell populations displayed significantly wider distributions in width **(Figure 4B)**. Once again, the *ΔmcdB* and *ΔmcdAB* mutant populations displayed a significant number of asymmetrical division events (~ 70 % of dividing cells) compared to wild-type or *ΔmcdA* **(Figure 4C)**. We propose that carboxysome mispositioning and aggregation at 30°C elicits cell division arrest, cell elongation and, when McdB is absent, asymmetric cell division; phenotypes that were largely masked when cells were grown at 40°C.

**Figure 4.**
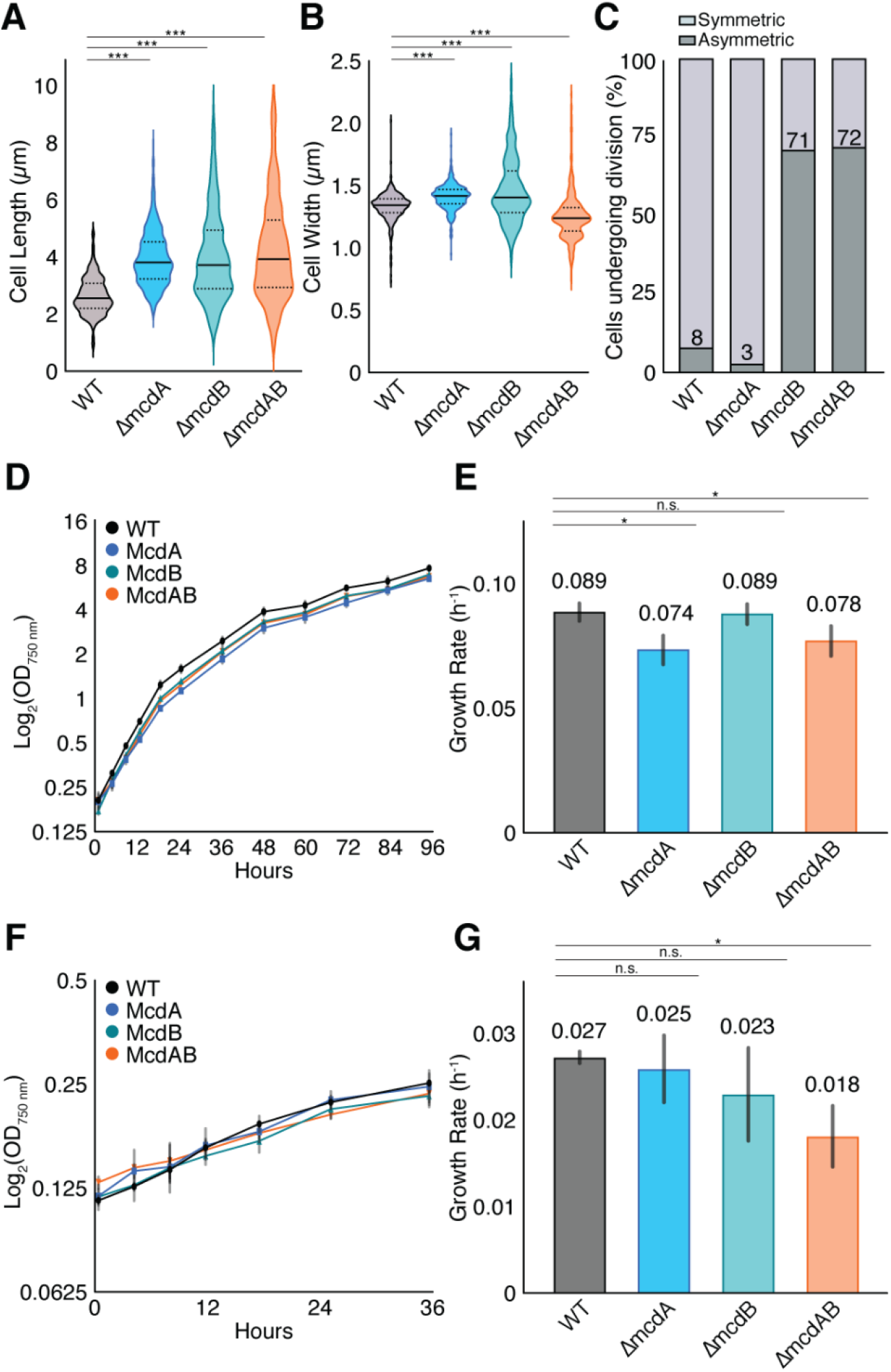
McdAB mutants display cell elongation and asymmetric cell division phenotypes at 30°C. (**A**) Cell lengths and (**B**) widths of WT population compared to McdAB system mutants. Cells were grown in 2% CO_2_. n = 440 cells of each strain. Statistical significance was based on a nonparametric Mann-Whitney test. *** = P < 0.001, ** = P < 0.01, * = P < 0.05, n.s. = not significant. (**C**) Percentage of cells undergoing symmetric (mid-cell; light grey bar) or asymmetric (non-mid-cell; dark grey bar) division. (**D**) Growth curves of all strains grown at 30°C in 2% CO_2_. (**E**) Quantification of mean exponential growth rate at 30°C in 2% CO_2_. (**F**) Growth curves of all strains grown at 30°C in 0.4% CO_2_. (**G**) Quantification of mean exponential growth rate at 30°C in 0.4% CO_2_. (**D-G**) Error bars represent the standard deviation from three independent biological replicates. Statistical significance was based on an unpaired t-test. *** = P < 0.001, ** = P < 0.01, * = P < 0.05, n.s. = not significant.

The changes in cell morphology suggested that the mutants may display slower growth rates, even when grown at high CO_2_. However, that was not the case. Under high CO_2_ conditions, no significant reduction in growth rate was observed for all three mutants when compared to wild-type **(Figure 4D-E)**. Even when grown in ambient CO_2_, reductions in growth rate of the mutants were minor **(Figure 4F-G)**.

Overall, we find that at 30°C, carboxysomes are mispositioned in the *ΔmcdA*, *ΔmcdB*, and *ΔmcdAB* mutants; with drastic increases in carboxysome spacing due to aggregation, resulting in fewer carboxysome foci per cell. We unveiled a cell elongation phenotype for all three mutant populations, and strikingly, we also found an asymmetric cell division phenotype that was exclusive to cells lacking McdB; a phenotype also found at 40°C but exacerbated at 30°C **(compare Panel C in Figures 2 and 4)**. Despite these carboxysome aggregation and cell-morphology phenotypes, only minor decreases in growth rate were observed. We propose that carboxysome aggregation decreases the carbon-fixing activity of encapsulated RuBisCO, which triggers an organic carbon-limitation response - cell division arrest, elongation, and asymmetric cell division.

### *S. elongatus* elongates at 20°C and this mode of growth is slower in Mcd system mutants

A ten degree drop in growth temperature unveiled the physiological consequences of carboxysome mispositioning at 30°C, which were largely masked at 40°C. The catalytic rate of carboxysome-encapsulated RuBisCO is often considered the bottle neck of photosynthesis because the enzyme is inefficient (28) and temperature dependent (20). The light reactions of photosynthesis, however, are considered temperature independent (20, 29). We therefore decided to study the effects of carboxysome mispositioning when cells were grown at 20°C, which is within the environmentally relevant temperature range - *S. elongatus* PCC 7942 was originally isolated from a fresh water source in the San Francisco bay area, with an annual temperature range of 8°C to 25°C (30).

Unexpectedly, we found that even wild-type *S. elongatus* undergoes extreme cell elongation when grown at 20°C **(Figure 5A)**. *S. elongatus* has been previously studied at this temperature (31), but to our knowledge, this is the first time cell morphology has been directly observed. Despite the dramatically long cell lengths, carboxysomes were still robustly aligned down the entire longitudinal axis of the cell **(Figure 5A)**. All three mutants were also elongated, but to a lesser extent, and carboxysomes were clearly mispositioned **(Figure 5B-D)**.

**Figure 5.**
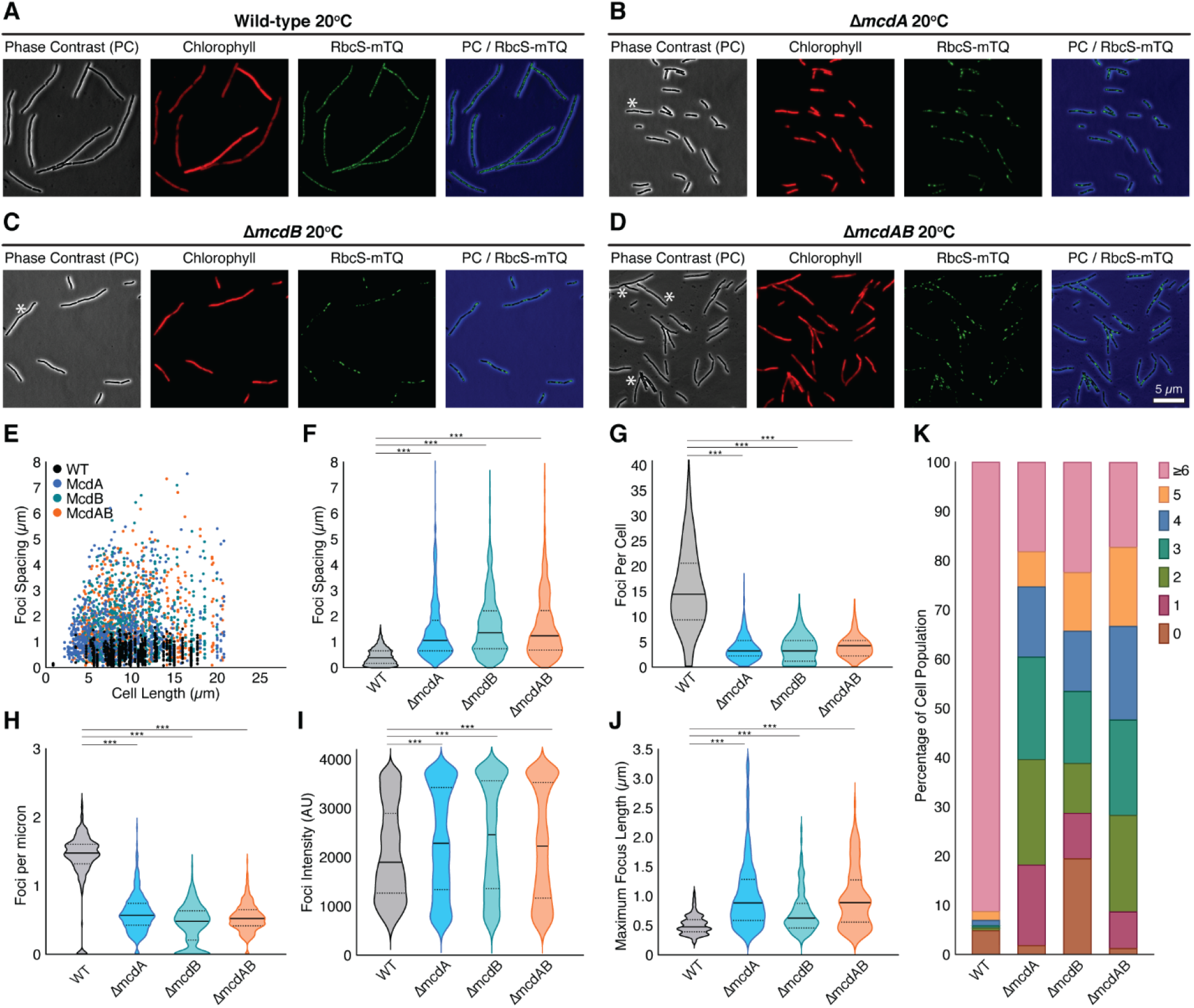
At 20°C wild-type and McdAB system mutants elongate, but only the mutants display few and mispositioned carboxysome aggregates. (**A-D**) Microscopy images of the specified cell strains grown in 2% CO_2_. Phase contrast (black), chlorophyll (red), and carboxysomes (green). Phase Contrast (PC) channel is blue in the merge for better contrast. (**E**) Spacing between carboxysome foci as a function of cell length. (**F**) Spacing between carboxysome foci in the same cell. (**G**) Carboxysome foci number per cell. (**H**) Number of carboxysome foci per unit cell length for each strain. (**I**) Carboxysome peak foci intensity for each cell strain. AU = Arbitrary Units. (**J**) Quantification of the maximum length across carboxysome foci. (**F-J**) Solid bars represent the median and dashed lines demarcate the 95% confidence interval. Statistical significance was based on a nonparametric Mann-Whitney test. *** = P < 0.001, ** = P < 0.01, * = P < 0.05, n.s. = not significant. (**K**) Population percentages of cells with the specified number of carboxysome foci. n ≥ 1000 carboxysomes from 440 cells of each strain.

Wild-type showed the same uniform carboxysome spacing distance (0.50 ± 0.30 μm) even in cells as along as 20 microns **(Figure 5E)**. All three mutants, on the other hand, displayed dramatically increased carboxysome spacing, and variability in spacing, as cell length increased **(Figure 5F)**. Since all cell types dramatically elongated at 20°C, the loss of carboxysome positioning resulted in a massive reduction in the number of carboxysome foci per cell **(Figure 5G)**, and per unit cell length **(Figure 5H)**. Carboxysome foci in all three mutants were significantly larger than that of wild-type, suggesting carboxysome aggregation **(Figure 5I-J)**. While ~ 90% of wild-type cells (n=388) had six or more foci, ~ 80% of all mutant populations had less than five **(Figure 5K)**. The data implicates carboxysome positioning by the McdAB system as a requirement for maximizing carbon-fixation in cells that are elongated when grown at cooler but environmentally relevant temperatures.

We quantified cell length at 20°C and found that the median length of wild-type cells was significantly longer than all three mutant populations **(Figure 6A)**. The *ΔmcdA* cells were similar in width to that of wild-type, while *ΔmcdB* and *ΔmcdAB* cell populations were notably thinner **(Figure 6B)**. Consistent with the wide distributions in cell length **(Figure 6A)**, a significant fraction of division events at this growth temperature were asymmetric across all cell populations. But the frequency of asymmetric division events was still highest in the strains lacking McdB **(Figure 6C)**.

**Figure 6.**
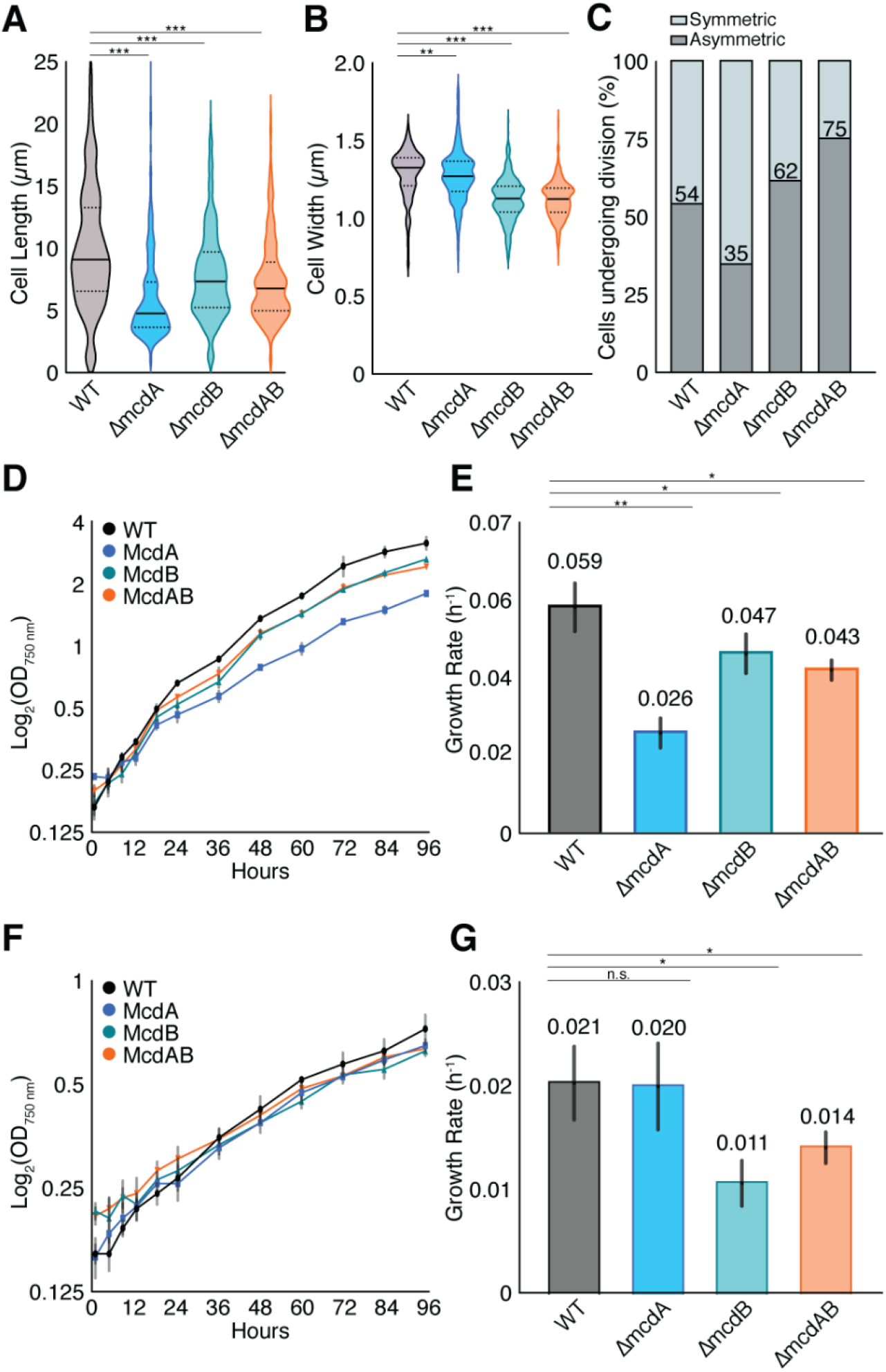
At 20°C wild-type and McdAB mutants elongate and undergo asymmetric cell division, but only mutants show slowed growth rates. (**A**) Cell lengths and (**B**) widths of WT population compared to McdAB system mutants. Cells were grown in 2% CO_2_. n = 440 cells of each strain. Statistical significance was based on a nonparametric Mann-Whitney test. *** = P < 0.001, ** = P < 0.01, * = P < 0.05, n.s. = not significant. (**C**) Percentage of cells undergoing symmetric (mid-cell; light grey bar) or asymmetric (non-mid-cell; dark grey bar) division. (**D**) Growth curves of all strains grown at 20°C in 2% CO_2_. (**E**) Quantification of mean exponential growth rate at 20°C in 2% CO_2_. (**F**) Growth curves of all strains grown at 20°C in 0.4% CO_2_. (**G**) Quantification of mean exponential growth rate at 20°C in 0.4% CO_2_. (**D-G**) Error bars represent the standard deviation from three independent biological replicates. Statistical significance was based on an unpaired t-test. *** = P < 0.001, ** = P < 0.01, * = P < 0.05, n.s. = not significant.

When growth rates were assayed at high CO_2_ **(Figure 6D-E)** or ambient CO_2_ **(Figure 6F-G)**, we finally found statistically significant reductions in growth rate for all three mutants when compared to wild-type. However, the reductions in growth rate depended on whether McdA or McdB was absent, and whether the cells were grown with high or ambient CO_2_. First, the *ΔmcdA* growth rate was 2-times slower than wild-type at high CO_2_ **(Figure 6E)**. But when grown in ambient CO_2_, the *ΔmcdA* growth rate was similar to wild-type **(Figure 6G)**. It is known that carboxysome quantity significantly increases when *S. elongatus* is grown in ambient CO_2_ (32). Therefore, our data suggest that with ambient CO_2_, the greater number of carboxysomes compensates for a loss in their positioning. Alternatively, a decrease in growth rate for *ΔmcdB* and *ΔmcdAB* was found with high- or ambient-levels of CO_2_ **(Figure 6E and G)**. The data once again shows that McdB plays a key role in carboxysome function outside of its role in positioning with McdA.

Overall, we identified a cell elongation phenotype in wild-type *S. elongatus* cells that occurs when grown at colder, but environmentally relevant temperatures. All three Mcd system mutants elongated to a lesser extent and have aggregated carboxysomes, which resulted in drastically fewer carboxysome foci per cell compared to wild-type. Significant decreases in growth rate were observed, even with high CO_2_ present. The findings suggest that in Mcd mutant or wild-type cells, colder growth temperatures decrease the enzyme kinetics of RuBisCO to the point where cell division arrest, elongation, and asymmetric cell division is triggered in response to carbon-limitation. At 30°C, this response was only found in the Mcd mutants likely due to reduced carbon-fixation efficiency resulting from carboxysome aggregation. At 40°C, the enzyme kinetics of RuBisCO are fast enough to compensate for carboxysome aggregation, which would explain why Mcd mutants largely displayed cell morphologies and growth rates similar to that of wild-type at higher growth temperatures.

### McdAB mutants have significantly decreased levels of intracellular RuBisCO

RuBisCO is the sole enzyme providing organic carbon to cellular biomass production and the phototrophic growth of *S. elongatus*. Therefore, we set out to determine if carboxysome mispositioning and the changes in cell morphology of McdAB mutants at 30°C correlated with altered cellular levels of RuBisCO. Immunoblot analysis against the Large subunit of RuBisCO (RbcL) showed that RuBisCO content is roughly half in all three mutants compared to wild-type cells **(Figure 7A-B)**. RbcL content from cell cultures was normalized based on Atpβ quantity from immunoblot analysis as shown previously (33, 34). The data suggest that although the mutants house massive carboxysome aggregates that are much brighter than those uniformly positioned in wild-type cells **(Supplemental Figure S2)**, the loss of a functioning McdAB system results in a reduction in the total cellular content of RuBisCO.

**Figure 7.**
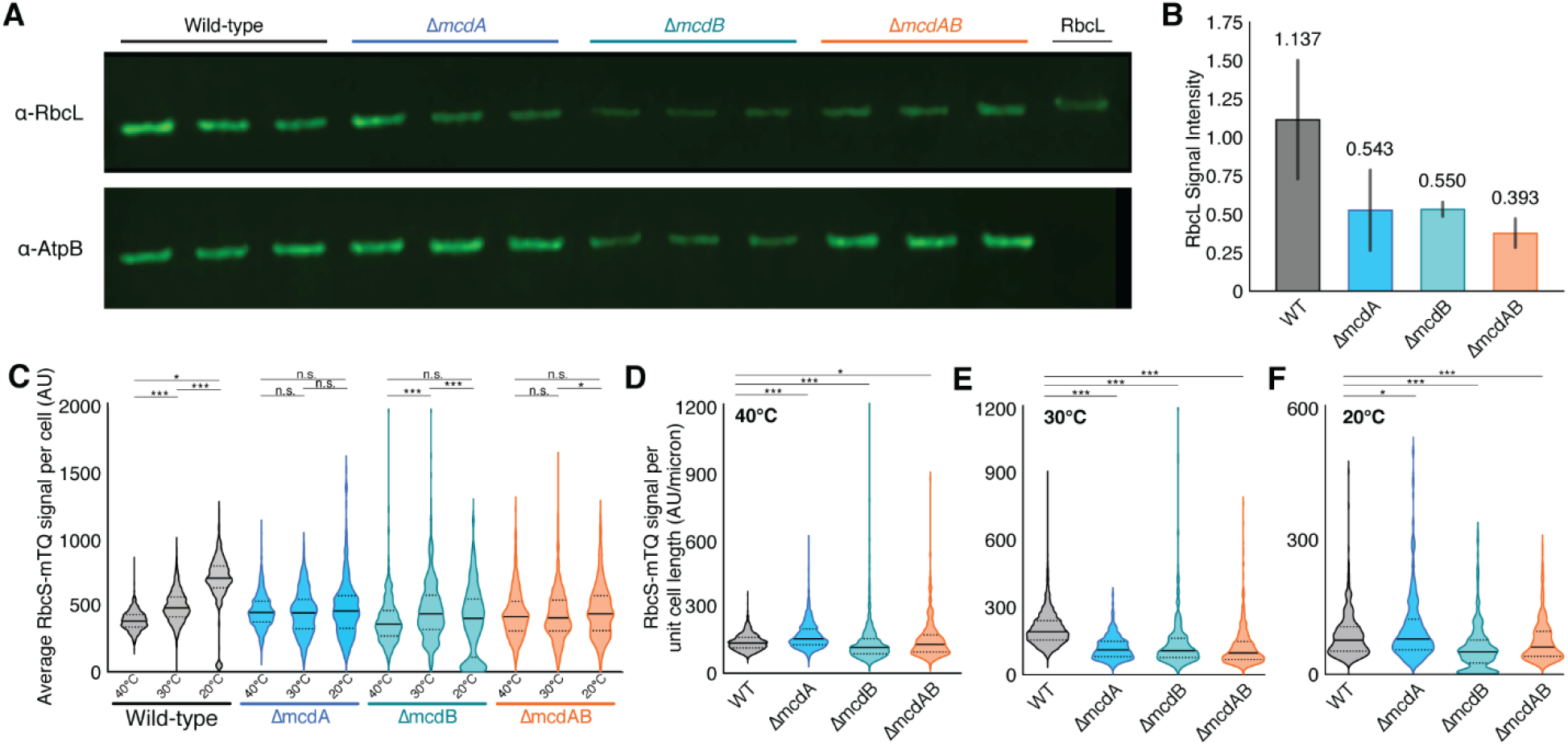
McdAB mutants have decreased cellular levels of RuBisCO. (**A**) Western Blot analysis of RbcL signal (top) and Atpß (bottom). Wild-type (lanes 1-3), *ΔmcdA* (lanes 4-6), Δ*mcdB* (lanes 7-9), and Δ*mcdAB* (lanes 10-12), purified RbcL control (lane 13). (**B**) Quantification of immunoblot signal for the cell type specified normalized against Atpβ signal. Cells were grown at 30°C in 2% CO_2_. Error bars represent the standard deviation from three independent biological replicates. (**C**) Quantification of the average RbcS-mTQ intensity per cell in each strain at the specified temperatures. (**D-F**) Quantification of the average RbcS-mTQ intensity normalized to cell length in each strain at the specified temperatures. n ≥ 338 per cell population. Statistical significance was based on an unpaired t-test. *** = P < 0.001, ** = P < 0.01, * = P < 0.05, n.s. = not significant.

We continued our quantification of cellular RuBisCO content using the RbcS-mTQ fluorescence signal. When quantifying the average RbcS-mTQ intensity per cell, only wild-type seemed to increase RuBisCO content in response to decreasing growth temperature **(Figure 7C)**. This finding is consistent with previous studies showing that RuBisCO abundance in phytoplankton populations (35), including cyanobacterial species (44), inversely correlates with growth temperature. We then normalized the RbcS-mTQ signal to cell length to account for the differences in cell morphology with Mcd system mutants and varying growth temperatures. At 40°C, RuBisCO content was similar across all cell populations, albeit more variable in the mutants **(Figure 7D)**. At 30°C, all three mutants had roughly 2-fold less RuBisCO compared to that of wild type **(Figure 7E)**; a decrease that is consistent with our immunoblot quantification at 32°C **(see Figure 7B)**. The data confirm that the immunoblot analysis using Atpβ standardization provided a valid measure of cellular RuBisCO abundance, and that the RbcS-mTQ intensity measurements *in vivo* are a reasonable proxy for RuBisCO content. Finally, at 20°C, all cell populations including wild-type had the lowest levels of RuBisCO content when normalized to cell length **(Figure 7F)** compared to that at higher temperatures **(Figure 7D-E)**. Also, cells without McdB had almost 2-fold less RuBisCO compare to wild-type or *ΔmcdA* cells **(Figure 7F)**. Overall, our findings suggest that carboxysome mispositioning results in decreased levels of cellular RuBisCO, and unveiled a critical but currently unknown role for McdB in carboxysome integrity and carbon-fixing function.

## DISCUSSION

We recently identified the McdAB system responsible for the equidistant positioning of carboxysomes in the rod-shaped cyanobacterium *S. elongatus* PCC 7942 (16). We also recently found that McdAB systems are widespread among β-cyanobacteria (17). These findings suggest important and widespread, but currently unknown, physiological roles for carboxysome positioning. Here, we show for the first time that a failure to distribute carboxysomes leads to a number of temperature-dependent changes in cell physiology: decrease in cell growth rate, cell division arrest, cell elongation, asymmetric cell division, and a significant reduction in cellular levels of RuBisCO.

At the three temperatures tested (20, 30, and 40°C), all three mutant cell-types housed few and irregularly spaced carboxysome aggregates, compared to wild-type cells with uniformly-spaced and -sized carboxysome foci. The data here provides quantitative support for our previous findings that the McdAB system equally distributes carboxysomes to opposite sides of the cell to ensure their inheritance following cell division (2, 16), akin to ParA-based plasmid partition systems in bacteria (4). But in addition, and particularly important for protein-based cargoes, the McdAB system serves to prevent carboxysome aggregation. This ‘anti-aggregation’ activity serves as a homeostasis mechanism that regulates carboxysome size, number, composition, positioning, and ultimately, its carbon-fixing function in the cell. It has recently been found that the pyrenoid, the functional analog of the carboxysome in the chloroplasts of the model alga *Chlamydomonas*, is also spatially regulated and this activity affects carbon-fixation (36). It remains to be seen if other BMCs also use spatial regulation mechanisms to avoid aggregation during their self-assembly to optimize enzymatic efficiency.

### Physiological defects associated with carboxysome mispositioning are temperature-dependent

Despite the drastic mispositioning and aggregation of carboxysomes in the *mcdA*, *mcdB*, and *mcdAB* deletion strains of *S. elongatus*, only minor changes in cell morphology and growth rate were found when compared to wild-type at the most optimal growth temperature used in this study (40°C) **(Figures 2)**. At 30°C however, all three mutants displayed filamentous growth, and a significant fraction of cell division events in the mutant populations lacking McdB were asymmetric **(Figure 4)**. Despite these changes to cell morphology, the growth rates were similar to that of wild-type; even in ambient CO_2_. At our coldest but environmentally relevant growth temperature of 20°C, we found that all cell populations, even wild-type, underwent filamentous growth and asymmetric cell division **(Figure 6)**. Wild-type cells at 20°C were as much as 5 to 10-times longer than the median length when grown at 30°C or 40°C. Even in these extremely elongated cells, the McdAB system robustly distributed carboxysomes down the entire cell length **(Figure 5A)**. This mode of filamentous growth was slower in all Mcd system mutants, resulting in significantly shorter filaments compared to wild-type **(Figure 6)**. Overall, we find temperature-dependent physiological defects associated with carboxysome mispositioning and aggregation; phenotypes that were masked at high growth rates and temperatures typically used in the lab, and unveiled at lower but environmentally relevant temperatures.

### Cell division arrest, filamentation, and asymmetric cell division – A stress response to carbon-limitation?

Many bacteria can change shape in response to growth conditions (37). *E. coli* and *B. subtilis* produce larger cells under nutrient-rich conditions and smaller cells under nutrient-limited conditions (38, 39). *Pseudomonas aeruginosa* elongates to enhance nutrient uptake during carbon and nitrogen starvation (40); *Caulobacter crescentus* cells increase cell area in response to phosphate starvation (41); and abundant human gut species such as *Bacteriodes thetaiotaomicron* elongate under sugar-limited conditions (42). Our findings suggest that *S. elongatus* filamentation and asymmetric cell division are responses triggered by organic carbon-limitation, resulting from reduced carbon-fixation activity of RuBisCO in this obligate photoautotroph. Cell filamentation (i) prevents the birth of cells devoid of carboxysomes, (ii) gives the cell an opportunity to increase carbon-fixation by producing more RuBisCO as a larger aggregate, or by nucleating the formation of additional carboxysomes *de novo* elsewhere along the elongated cell and (iii) the increased cell-surface area would increase the light-harvesting capability for photochemistry.

A carbon-limitation response parsimoniously explains the two triggers identified here: (1) carboxysome aggregation and (2) colder growth temperatures. What is the mechanism behind the carboxysome aggregation trigger? All Mcd system mutants displayed significant reductions in their total cellular RuBisCO content **(Figure 7)**. Therefore, carboxysome aggregation could trigger the carbon-limitation response simply due to lower levels of RuBisCO as a result of losses in carboxysome integrity and leakage, enzyme stability, and/or degradation. Alternatively, wild-type cells have carboxysomes of a homogenous size, and as cells grow, so do the number of carboxysomes and their net surface area. The net surface area of the carboxysome shell, where RuBisCO substrates and products interface with the cytoplasm, significantly decreases due to carboxysome aggregation. As a result, substrate and product permeability would suffer, which could lead to decreased carbon-fixation rates by encapsulated RuBisCO. It is therefore also possible that the carbon-limitation response is triggered by decreased RuBisCO activity due to decreases in the effective surface area of aggregated carboxysomes. These two proposals are not necessarily mutually exclusive from one another. Consistent with these proposals, it was recently found in Synechococcus sp. PCC 7002 that mispositioning of inactivated carboxysomes towards the cell poles is strongly correlated with their rapid degradation (35).

We propose this adaptive response to carbon-limitation is also triggered by low temperature growth, even in wild-type *S. elongatus*. At high temperature (i.e. 40°C), RuBisCO activity is high (43) and sufficient to compensate for carboxysome mispositioning and RuBisCO aggregation in Mcd system mutants. Therefore, the carbon-limitation response is not triggered. At 30°C, RuBisCO activity is still high enough to prevent a carbon-limitation response in wild-type cells, but not in Mcd mutants with few and aggregated carboxysomes. As a result, cell filamentation is triggered only in the Mcd mutants. At 20°C, RuBisCO activity has reduced to a level that, even in wild-type cells, triggers the carbon-limitation response. Mcd mutants are starved for carbon to the point where growth is slowed, resulting in shorter filaments compared to wild-type.

### Spatial regulation of carboxysomes influences RuBisCO activity and abundance

Psychrophilic phytoplankton species deal with low temperatures, well below the thermal optimum of most enzymes, by increasing the abundance of RuBisCO (35). It has recently been found that low temperature growth (18 to 22°C) results in increased RuBisCO levels in the marine cyanobacterium *Synechococcus* PCC 7002, compared to growth at 26°C or 30°C (44). This trend was also observed in natural phytoplankton assemblages across a wide latitudinal range (35). Consistently, we found here that the RbcS-mTQ signal within carboxysome foci, and within the entire cell, inversely correlated with decreasing growth temperatures.

Previous studies have also shown positive relationships between growth rate and RuBisCO abundance (35, 44–48). Elevated carbon fixation rates during blooms in polar regions have been shown to be associated with several-fold increases in RuBisCO. Twenty degrees Celsius is well below the thermal optimum for RuBisCO activity (49), and well below the optimal growth temperature of *S. elongatus*. We therefore conclude that increasing RuBisCO abundance and triggering cell elongation both represent acclimation responses that compensate for the decreased catalytic rate of RuBisCO at low-temperature growth. This response is further compounded when carboxysomes are mispositioned and aggregated in our Mcd mutants. It is important to note that filamentation occurred here under high CO_2_ – a condition whereby *S. elongatus* does not require the carbon-concentrating activity of carboxysomes for growth (50). Therefore, the evidence further supports that it is not CO_2_ substrate limitation causing filamentation, but rather RuBisCO activity that is significantly compromised when complexed in carboxysome aggregates or when cells are grown at colder temperatures. These findings shed light on the importance of the McdAB system in maximizing the carbon-fixation activity of RuBisCO by preventing aggregation during carboxysome self-assembly.

Overall, we show that carboxysomes are organized by the McdAB system in cyanobacteria and can appropriately respond to variability in RuBisCO abundance and activity along a wide temperature gradient. Our results suggest that carboxysome homeostasis provided by the McdAB system is part of an autotrophic growth strategy in elongated cells - RuBisCO abundance increases and is distributed into homogeneously-sized carboxysomes at low temperature to overcome the lower catalytic rates of this temperature-dependent enzyme (35).

### McdB plays a role in carboxysome function outside of its positioning with McdA

Regardless of growth temperature, Δ*mcdB* and Δ*mcdAB* mutants elicited stronger cell elongation and asymmetric division phenotypes compared to Δ*mcdA*. Our findings suggest that McdB plays a currently unknown but critical role in carboxysome function, outside of its role in positioning with McdA. Consistently, our recent bioinformatics analysis of McdAB systems across cyanobacteria identified numerous species with orphan McdBs, once again suggesting functional roles independent of McdA (17). We have also shown previously via bacterial two-hybrid analysis that McdA does not physically associate with any carboxysome component, while McdB directly interacts with a number of shell proteins (16). We have also recently found that purified McdB undergoes Liquid-Liquid Phase-Separation (LLPS) *in vitro* (17). This activity is intriguing given recent studies showing that both α- and β-carboxysomes, as well as the algal pyrenoid, have intrinsically-disordered proteins that from liquid-like condensates with RuBisCO (51–53). Collectively, these studies suggest that LLPS is a common feature underlying carboxysome biogenesis. It is intriguing to speculate that the LLPS activity of McdB and other carboxysome components are related and potentially influence each other.

### Cold growth temperature triggers elongation and asymmetric cell division

Across the bacterial world, complex systems maintain cell-size homeostasis. We find here that *S. elongatus* maintains cell-size homeostasis when grown at 30°C or 40°C, but undergoes filamentation when grown at the environmentally-relevant temperature of 20°C. A growing number of bacterial species are known to elongate into a filamentous morphology amidst environmental changes to promote survival. For example, *E. coli* cells become filamentous during infection (54, 55), and during DNA damage, the SOS response blocks cell division until damage has been repaired, resulting in elongation (56). Several other forms of stress have been shown to induce elongation, including host environment, antibiotics, nutrient access, pH, heat shock, and osmotic fluctuations (57–60). Bacteria can clearly exist in diverse morphological states, in part dictated by their environment (37, 61).

To our knowledge, we find here the first example of bacterial filamentation caused by cold-growth temperature. We propose colder temperatures trigger filamentation indirectly as a carbon-limitation response due to the reduced carbon-fixing activity of RuBisCO. Alternatively, transition into a filamentous morphology has been proposed to confer advantages, such as avoiding phagocytosis via the host immune response (54, 55, 57, 61) or to protect against predation in aquatic environments (58, 62–64). It is attractive to speculate that the growth temperature of 20°C coincides with the seasonal temperature during which predation occurs (63). That is, *S. elongatus* may use temperature as a cue to elongate so as to avoid planktivorous protists. Consistent with this possibility, previous studies have demonstrated that several freshwater bacteria, including *Caulobacter crescentus* (58), exhibit high phenotypic plasticity and can transition to a filamentous morphology – a transition that may be specifically triggered in the presence of a size-selective protistan predator (63, 65, 66). It is possible that some aquatic bacteria have evolved to undergo filamentation at temperatures that coincide with grazing season.

Several studies have characterized the mechanisms by which bacterial filamentation is induced, particularly under DNA damage or exposure to antibiotics (56). But how bacterial cells enter and exit these filamentous states to ensure survival during changes in environment remains poorly characterized and is a recent question of interest. We find here that *S. elongatus* forms filaments when grown at low temperature or when carboxysomes are mispositioned. When these filaments divide, it is asymmetric – forming a daughter cell of ‘normal’ length. It has been recently shown that *S. elongatus* can also be induced to filament under dim-light stress, and then divides asymmetrically to form daughter cells of the correct size when brought back into well-lit conditions (67). Somewhat related, chlorophyll fluorescence along the cell length of our Mcd mutants was much more heterogeneous, particularly at 20°C, compared to wild-type **(see Figure 5A-D)**. It should be noted that in addition to sensing light and inorganic carbon substrates, *S. elongatus* responds to lowering temperatures by slowing and even turning off circadian rhythms; thereby down regulating cell growth, division, and metabolic flux (68, 69).

The Min system has been shown to play a role in the asymmetric cell division and daughter-cell sizing of filamentous *E. coli* (60), *V. parahaemolyticus* (70, 71), and *S. elongatus* (67). Recent findings suggest that division restoration at the poles of these filaments is regulated by a combination of Min oscillations, FtsZ levels and *terminus* segregation, resulting in daughter cells of the right length (72, 73). The mechanism by which filamentation and asymmetric division occurs in *S. elongatus* is an area of future research. Taken together, the conserved ability for various bacterial species to undergo filamentation and morphological recovery, some of which showing direct selective benefits, strongly suggests that this differentiation plays an important role in survival and proliferation.

## MATERIALS AND METHODS

### Construct designs

All constructs in this study were generated using Gibson Assembly (74) from synthetized dsDNA and verified by sequencing. Constructs contained flanking DNA that ranged from 500 to 1500 bp in length upstream and downstream of the targeted insertion site to promote homologous recombination into target genomic loci (75).

### Generation of Bacterial Strains

All *Synechococcus elongatus* PCC7942 transformations were performed as previously described (75). Plasmid constructs of *mcdA*, *mcdB*, and *mcdAB* deletions were created by replacing the respective coding sequences with a kanamycin resistance cassette. All fluorescent strains were transformed using plasmid pAH40, which contains a chloramphenicol resistance cassette and a second copy of the *rbcS* promoter and gene, attached at the 3’ end with the gene encoding for fluorescent protein mTurquoise2 (mTQ) and separated with a GSGSGS linker, inserted into neutral site 1. Transformed cells were plated on BG-11 agar. Single colonies were picked into 96-well plates containing 300 μL of BG-11 with 6.35 ug ml^−1^ kanamycin or 3.125 ug ml^−1^ chloramphenicol. Concentrations of both respective antibiotics were gradually increased to 12.5 μg ml^−1^. Cultures were verified for complete insertion via PCR and removed from antibiotics for experiments.

### Growth Conditions

Both wild-type and mutant *Synechococcus elongatus* PCC7942 strains were grown in 125 mL culture flasks (Corning) in 50 mL BG-11 (Sigma) medium buffered with 1 g L^−1^ HEPES to pH 8.3 shaken at 130 rpm or on BG-11 plates containing 1.5% (w/v) agar. All strains were maintained and grown under constant LED illumination of 100 μE m^−2^ s^−1^ at either 20°C, 30°C, or 40°C in 2% or 0.04% CO_2_ as specified. Cultures grown in 0.04% CO_2_ were grown in air with no additional CO_2_. Cultures were regularly diluted with fresh medium to maintain exponential growth phase for subsequent imaging and immunoblot analyses. For cloning, One Shot™ TOP10 Chemically Competent *E. coli* (ThermoFisher) were grown aerobically at 37° in Luria-Broth medium.

### Fluorescence microscopy

Microscopy was performed using exponentially growing cells at an OD of 0.4. Two milliliters of culture were spun down at 15,000 x g for 60 s, resuspended in 100 μL of BG-11. Five microliters were transferred to a square 1.5% agarose + BG-11 pad, which was then flipped onto a 35 mm cell culture dish with a #1.5 glass coverslip bottom (ManTek). All images were captured using a Nikon Eclipse Ti2 inverted microscope with a PlanApo Objective lens (100x, 1.45NA, oil immersion), with phase contrast transillumination, and with a SOLA LED light source for imaging chlorophyll fluorescence (Excitation: 560/40nm (540-580nm), Emission: 630/70nm (593-668nm), Dichroic Mirror: 585nm) and RbcS-mTQ fluorescence (Excitation: 436/20nm (426-446nm), Emission: 480/40nm (460-500nm), Dichroic Mirror: 455nm). Images were acquired using a Photometrics Prime 95B Back-illuminated sCMOS Camera. Image analysis was performed using Fiji (76) and the MicrobeJ plugin (77).

### Growth Curve Measurements

Cultures were inoculated at a starting OD_750_ of 0.1-0.2 with fresh BG-11. Cell growth was monitored at OD_750_ using a DS-11 Spectrophotometer (DENOVI) at the specified time points. Growth rates were calculated using the linear portion of the exponential phase of growth. Error bars represent the standard deviation from three biological replicates that were recorded from different culture flasks.

### MicrobeJ quantification

Multiple fields of view were taken for each cell strain using three channels: Phase contrast provided cell perimeters, and fluorescence microscopy provided chlorophyll autofluorescence and RbcS-mTQ intensities. These data were analyzed using MicrobeJ (77). In each cell-line, cell length detection was performed using the rod-shaped descriptor and thresholding set to 0.4μm < area < max, 0.71μm < width range < 2 μm, 0 μm < width variation < 0.2 μm, and 0 μm < angularity amplitude < 0.35 μm. Carboxysome detection was performed using the point function with a tolerance of 20 and an intensity minimum of 500. Associations, shape descriptors, profiles, and distances were recorded for each strain. All MicrobeJ quantification was also verified manually. Graphs and statistical analyses were generated with Graph Pad Prism.

### Immunoblot Analysis

Cells were lysed with a Qsonica sonication system (20 cycles - 30 s on, 10s off at 30% power) in 0.5 mL RuBisCO extraction buffer (50 mM EPPS at pH 8.1, 1% PVPP, 1 mM EDTA, 10 mM DTT, 0.1% Triton, and sigma protease inhibitor). Laemmli sample buffer (0.1 mL of 4x) was added to lysate prior to loading 10 μL on a 4%–12% Bis-Tris NuPAGE gel (Invitrogen). Gels were transferred onto a Mini-size polyvinylidene difluoride membrane (Bio-Rad) using a Trans-Blot Turbo system (Bio-Rad). The membrane was immunoprobed using rabbit polyclonal antisera against RbcL and the beta subunit of ATP synthase, AtpB (Agrisera) and then goat anti-rabbit IgG secondary antibody (LI-COR). Membrane signals were visualized and quantified at 600 nm using LI-COR Image Studio. For each sample, there were three replicates from the same cell lysate. We normalized RbcL signal using AtpB; a method previously used in Zhang *et al*., 2012 and specifically with *S. elongatus* PCC7942 in Sun *et al*., 2016 (33, 34).

**Table 1:** Strains used in this study

| Strain | Description | Source |
| --- | --- | --- |
| *Synechococcus elongatus*<br>PCC7942 | Wild Type | - |
| *Synechococcus elongatus*<br>PCC7942 | $\Delta mcdA$ | (16) |
| *Synechococcus elongatus*<br>PCC7942 | $\Delta mcdB$ | (16) |
| *Synechococcus elongatus*<br>PCC7942 | $\Delta mcdAB$ | (16) |
| *Synechococcus elongatus*<br>PCC7942 | *rbcS-mTQ* | Wild Type transformed with<br>pAH40 |
| *Synechococcus elongatus*<br>PCC7942 | $\Delta mcdA + rbcS-mTQ$ | $\Delta mcdA$ transformed with pAH40 |
| *Synechococcus elongatus*<br>PCC7942 | $\Delta mcdB + rbcS-mTQ$ | $\Delta mcdB$ transformed with pAH40 |
| *Synechococcus elongatus*<br>PCC7942 | $\Delta mcdAB + rbcS-mTQ$ | $\Delta mcdAB$ transformed with pAH40 |

## ACKNOWLEDGMENTS

We would like to thank Drs. Lyle Simmons, Vincent Denef, and Daniel Ducat for providing critical feedback in experimental direction and data interpretation. We would also like to thank Pusparanee Anne Hakim for experimental assistance and critical reading of the manuscript.

## Supplemental Figures

**Figure S1.**
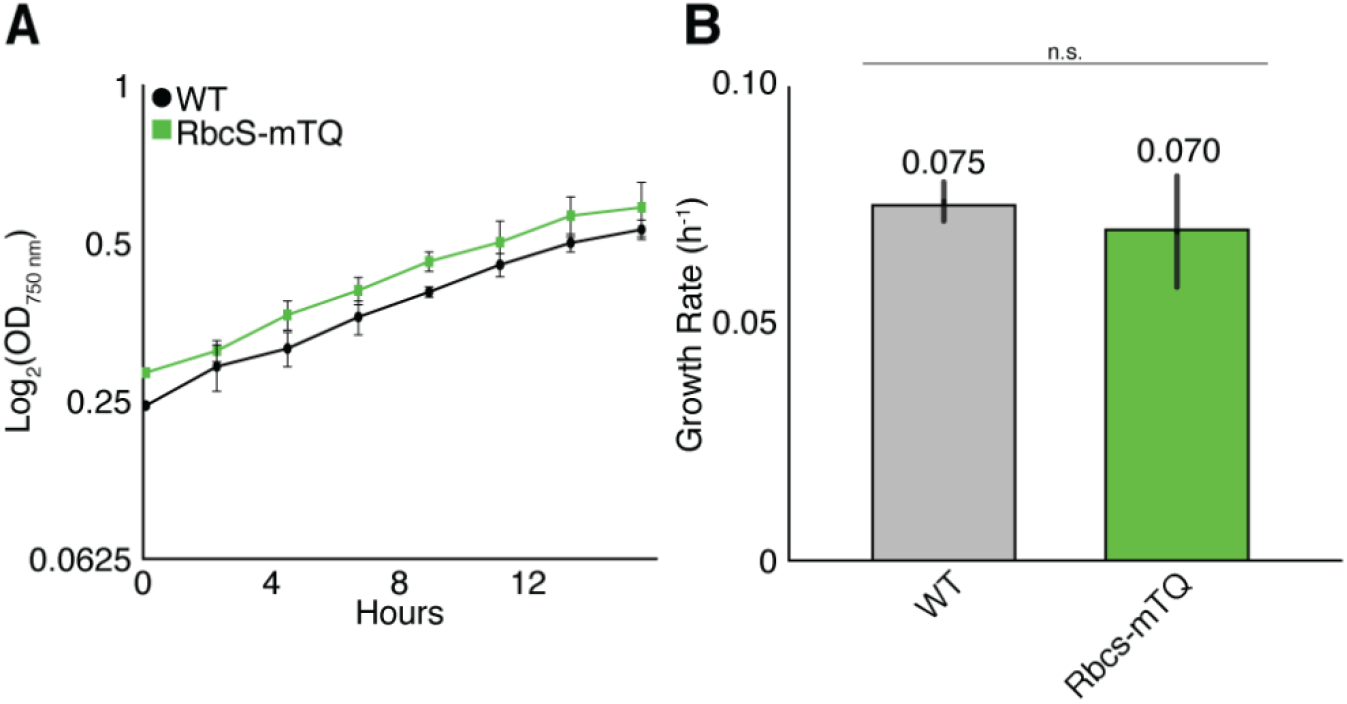
Presence of RbsC-mTQ carboxysome marker does not influence growth. (**A**) Growth curves of wild-type *S. elongatus* with and without RbcS-mTQ carboxysome marker. Cells were grown at 30°C in 2% CO_2_. (**B**) Comparison of growth rates. Error bars represent the standard deviation from three independent biological replicates. Unpaired t-test shows growth rates were not statistically different.

**Figure S2.**
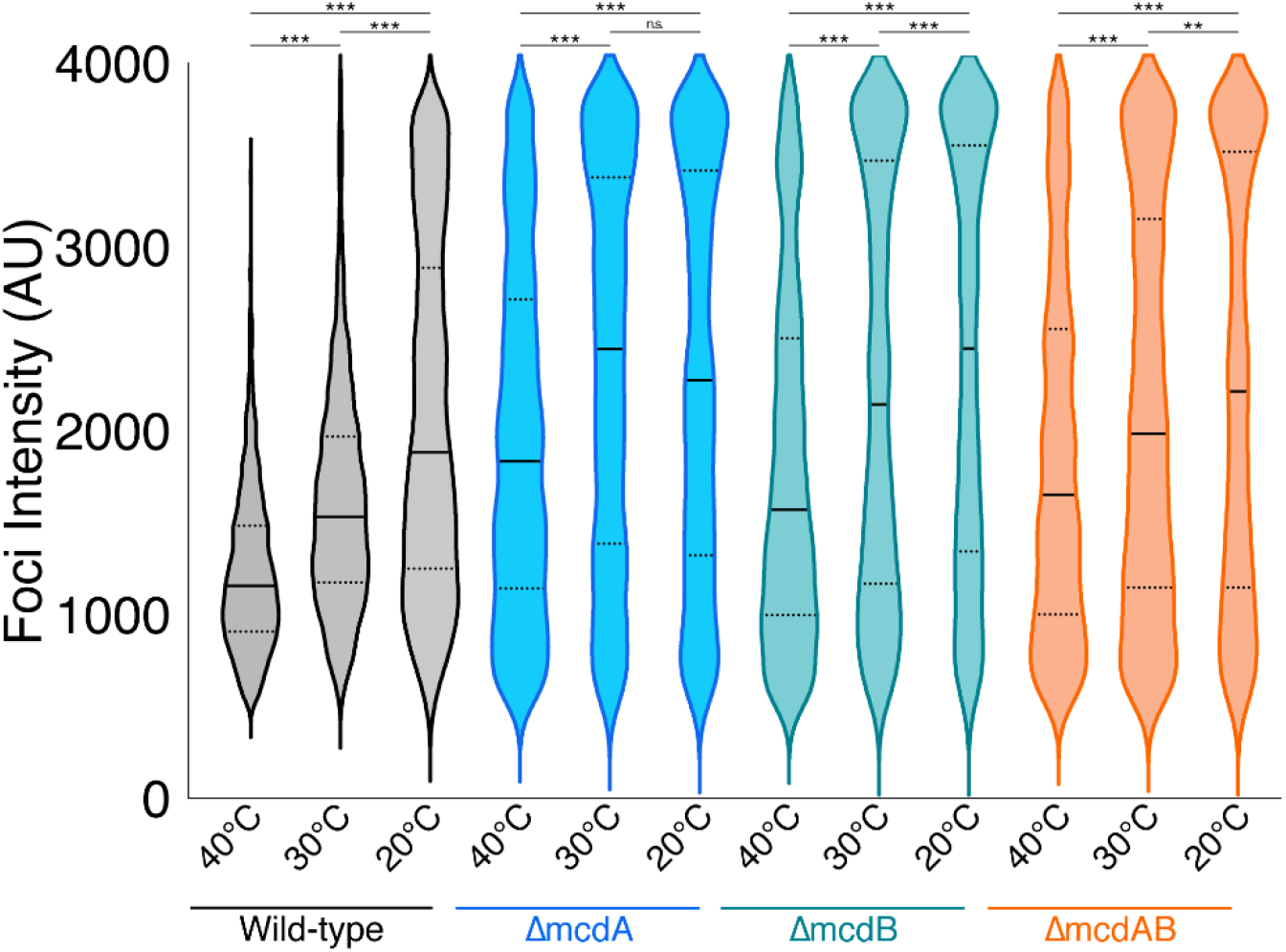
Comparison of the peak RbcS-mTQ foci intensities for each strain across the three growth temperatures tested in this study. AU = Arbitrary Units AU. n > 1000 carboxysomes per cell-line at each temperature. Solid bars represent the median and dashed lines demarcate the 95% confidence interval. Statistical significance was based on a nonparametric Mann-Whitney test. *** = P < 0.001, ** = P < 0.01, * = P < 0.05, n.s. = not significant.

